# Controlled dehydration, structural flexibility, and Gadolinium MRI contrast compound binding in human plasma glycoprotein afamin

**DOI:** 10.1101/673376

**Authors:** Andreas Naschberger, Pauline Juyoux, Jill von Velsen, Bernhard Rupp, Matthew W. Bowler

## Abstract

Afamin, a human blood plasma glycoprotein, a putative multi-functional transporter of hydrophobic molecules and a marker for metabolic syndrome, poses multiple challenges for crystallographic structure determination, both practically and in analysis of the models. Several hundred crystals were analysed, and unusual variability in cell volume and difficulty solving the structure despite a ~34% sequence identity with non-glycosylated human serum albumin indicated that the molecule exhibits variable and context-sensitive packing, despite greatly simplified glycosylation in insect cell expressed recombinant afamin. Controlled dehydration of the crystals was able to stabilise the orthorhombic crystal form reducing the number of molecules in the asymmetric unit from the monoclinic form and changing the conformational states of the protein. An iterative strategy, using fully automatic experiments available on MASSIF-1, was used to quickly determine the optimal protocol to achieve the phase transition that should be readily applicable to many types of sample. The study also highlights the drawback of using a single crystallographic structure model for computation modelling purposes given that conformational state of the binding sites and electron density in the binding site, likely resulting from PEGs, greatly varies between models. This also holds for the analysis of unspecific low-affinity ligands, where often a variety fragments with similar uncertainty can be modelled, inviting interpretative bias. As a promiscuous transporter, afamin also seems to bind Gadoteridol, a magnetic resonance imaging contrast compound, in at least two sites. One pair of Gadoteridol molecules is located near the human albumin Sudlow-site, and a second Gadoteridol at an intermolecular site in proximity of domain IA. The data from the co-crystals provide an opportunity to evaluate modern metrics of data quality in the context of the information that can be gleaned from data sets that would be abandoned on classical measures.

**Synopsis:** Controlled dehydration experiments have revealed a new crystal form of afamin, a human blood plasma glycoprotein and transporter of hydrophobic molecules. The comparison shows substantial molecular plasticity and amplifies the necessity to examine multiple crystal forms and to refine multiple models, while at the same time the new structure cautions against interpretation of fatty acid ligand density in crystals derived from PEG as major precipitants. An isomorphic low-resolution structure model suggests that afamin is capable of transporting Gadolinium-DO3A, a magnetic resonance imaging compound.

## 1.0 Introduction

Afamin (AFM) is a blood plasma glycoprotein and a member of the albumin gene family, reported as a multi-functional transporter of hydrophobic molecules such as vitamin E (Voegele *et al.*, 2002) and a potential binding partner for wnt signalling proteins (Mihara *et al.*, 2016). Afamin is a biomarker for metabolic syndrome, where elevated afamin levels are associated with all major parameters for metabolic syndrome such as high blood glucose, dyslipemia, obesity and high blood pressure as well as preeclampsia and ovarian cancer (Dieplinger *et al.*, 2009, Kronenberg *et al.*, 2014, Seeber *et al.*, 2014). A potential role of afamin in glucose metabolism in papillary thyroid carcinoma has been reported (Shen *et al.*, 2016). Afamin was shown *in vitro* to form a 1:1 complex with most of the 19 Wnt proteins (Mihara *et al.*, 2016), maintaining biological activity for Wnt3 and Wnt3a, and partial activity for Wnt1, Wnt7a, Wnt7b, Wnt8a, Wnt9b and Wnt10b, which play a crucial role in cell differentiation during embryogenesis and are involved in the development of various diseases including cancer (Nusse & Varmus, 2012, Nile & Hannoush, 2019). A compelling computational model of lipid-bound afamin in a monoclinic crystal form (PDB entry 5OKL) docked with a homology model of Wnt3A indicated that afamin can capture the acyl chain of palmitoylated Wnt3A serine 209 in a deep hydrophobic pocket (Naschberger *et al.*, 2017).

The preparation and crystallization of afamin has posed significant challenges and is non-trivial (Altamirano *et al.*, 2018). In contrast to human serum albumin (HSA), which is exported from the liver as a non-glycosylated chain, human afamin *in vivo* is highly and variably enzymatically glycosylated. Neither complex glycosylation nor possible unspecific binding of lipid components from the expression media bode well for crystallization. Only AFM purified from the Sf21 baculoviral system as described in §2.1 yielded crystals, and out of several hundred mounted crystals, only few data sets could be successfully processed (Naschberger *et al.*, 2017), revealing a large variability in cell dimensions and symmetry.

Troublesome crystals can sometimes be improved by dehydration. The reduction in the mole fraction of water surrounding crystals of macromolecules, either by changing the components of the mother liquor or using specific humidity control devices, can induce phase transitions (Heras & Martin, 2005, Newman, 2006, Russo Krauss *et al.*, 2012). These transitions can lead to an increase in the order within the crystal lattice, result in an increase in diffraction quality, or provide other beneficial changes such as an increase in symmetry or decrease in anisotropy (Bowler *et al.*, 2006, Cramer *et al.*, 2000, Hu *et al.*, 2011, Kadlec *et al.*, 2011, Raj *et al.*, 2017, Zerrad *et al.*, 2010, Scherer *et al.*, 2014). There are many examples of spectacular increases in diffraction quality where initially very poor data yield interpretable maps. Often small changes can have similarly decisive beneficial results. Here we describe how controlled dehydration of afamin crystals was able to stabilise a new orthorhombic crystal form increasing the resolution, inducing a change in symmetry and increasing the number of intermolecular contacts leading to the a more complete model. An iterative process using automated protocols on MASSIF-1 (Svensson *et al.*, 2015) to combine running automated dehydration experiments (Bowler, Mueller, *et al.*, 2015) with multi-crystal data collection (Svensson *et al.*, 2018) allowed the rapid determination of optimised conditions and selection of the best data set.

Human serum albumin is a promiscuous drug transporter known to transport paramagnetic magnetic resonance imaging (MRI) contrast enhancement agents (Fasano *et al.*, 2005). The structural similarity of afamin to HSA (Naschberger *et al.*, 2017) suggests that afamin might also be able to transport cargo in the blood stream. Tissue specific transport by Afamin could greatly enhance resolution of MRI imaging in blood-flow studies, for example (Schultz *et al,* 1999). As no crystal structure of albumin with a contrast agent is yet available, we selected Gadoteridol (Gd-DO3A, Figure 1), which is used in medical applications as a MRI contrast agent (Caravan, 2009) and in crystallography as a lanthanide phasing compound (Girard *et al.*, 2003). We were able to identify three likely Gd binding sites in a similar orthorhombic, low-resolution electron density map of AFM by co-crystallization with Gd-DO3A. In view of marginal quality of data and an incomplete model, details of Gd-DO3A binding remain tentative.

**Figure 1.**
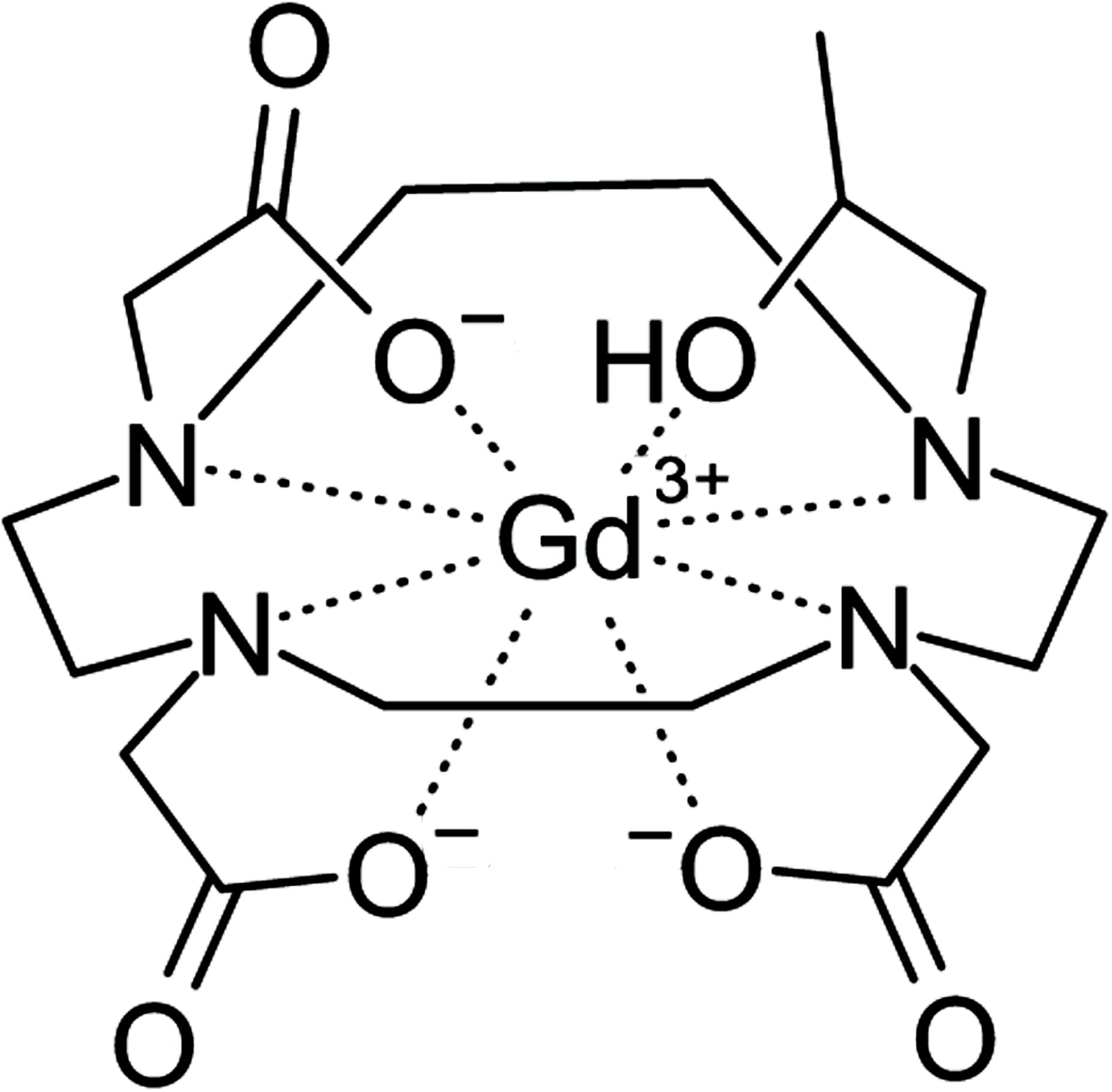
Structure of paramagnetic MRI contrast agent gadoteridol, Gd-DO3A.

The availability of 4 independent AFM structure models from two different crystal forms provides insight into the flexibility and dynamics of the molecule, which points towards a significant conformational adaptability when binding to hydrophobic molecules, metal chelates, or proteins with hydrophobic acylation as such as Wnt, supporting the function of afamin as a promiscuous human plasma transporter. The palmitoleic acid, present in Wnt proteins and identified by lipid analysis in afamin, could plausibly be modelled into electron density in a deep central hydrophobic cavity present only when the molecule adopted the most open conformation (Naschberger *et al.*, 2017). Modern structural biology beamlines now allow the collection of vast amounts of data, the option of refining multiple models should not be ignored.

## 2.0 Experimental details

### 2.1 Protein expression and purification

The challenges in preparing AFM (Uniprot entry P43652, AFAM_HUMAN) have been described (Altamirano *et al.*, 2018) and the successful expression of AFM in the *Spodoptera frugiperda* (Sf) 21 insect cell line (Emery, 1991) and subsequent purification of the glycoprotein for structural studies have been reported (Naschberger *et al.*, 2017).

### 2.2 Crystallisation

#### Native AFM crystals

Initial crystals were obtained at 291 K via the sitting drop method in a 96-well plate format (SwissSci 30926) set up with a Phoenix robot (Art Robbin Instruments, Sunnyvale, CA) using the PEG wizard screen (30 % PEG 1000, 200 mM CH_3_COONH_4_). Crystals were optimized using streak seeding, while 6-aminohexanoic acid (ACA) was added to a final concentration of 3% (w/v) in a hanging drop vapour-diffusion setup (VDX ™ plate Hampton HR3-142). 1.5 µL droplets of AFM stock solution (5 mg/ml SEC purified) were mixed with 1.5 µL of reagent (30% PEG1000, 170 mM CH_3_COONH_4_, 3% ACA) followed by micro-seeding immediately after pipetting with no pre-equilibration of the drop. The actual crystallization drop pH of 6.5 was determined by measurement. Different seed dilutions were obtained by loading a cat-whisker once with crushed crystals followed by sequential streak seeding into six different drops.

#### AFM-Gd-DO3A co-crystals

To obtain crystals of afamin containing gadoteridol (Gd-DO3A) the protein was incubated with 10 mM Gd-DO3A (NatXRay, Grenoble, France) and immediately sent for high throughput screening at the HTX laboratory (EMBL, Grenoble). Initial conditions were obtained in 25% PEG 4000, 0.2 M ammonium sulphate and 0.1 M sodium acetate pH 4.6. These conditions were refined with crystals appearing in 23 to 28% PEG 4000, 0.2 M ammonium sulphate and 0.1 M sodium acetate pH 5.5. All crystal were harvested by laser photoablation and cryo-cooled using the CrystalDirect Robot (Zander *et al.*, 2016, Pellegrini *et al.*, 2011).

### 2.3 Data collection

Prior to dehydration experiments, 348 crystals were screened (Figure 2) on the fully autonomous ESRF beamline MASSIF-1 (Bowler, Nurizzo, *et al.*, 2015, Nurizzo *et al.*, 2016) at a fixed wavelength of 0.9660 Å. Data were processed by the EDNA automated data processing pipeline (Monaco *et al.*, 2013) employing XDS and XSCALE (Kabsch, 2010) and pointless/aimless/ctruncate from the CCP4 program suite (Winn *et al.*, 2011) and xtriage from the phenix suite (Adams *et al.*, 2011). Strategy calculations accounted for flux and crystal volume in the collection parameter prediction for complete data sets and in the case of the monoclinic *mP* lattice, the symmetry was pre-set (Svensson *et al.*, 2015). Large scale screening was required as a wide variety of crystal volumes was observed with very few diffracting to high resolution (Figure 2).

**Figure 2.**
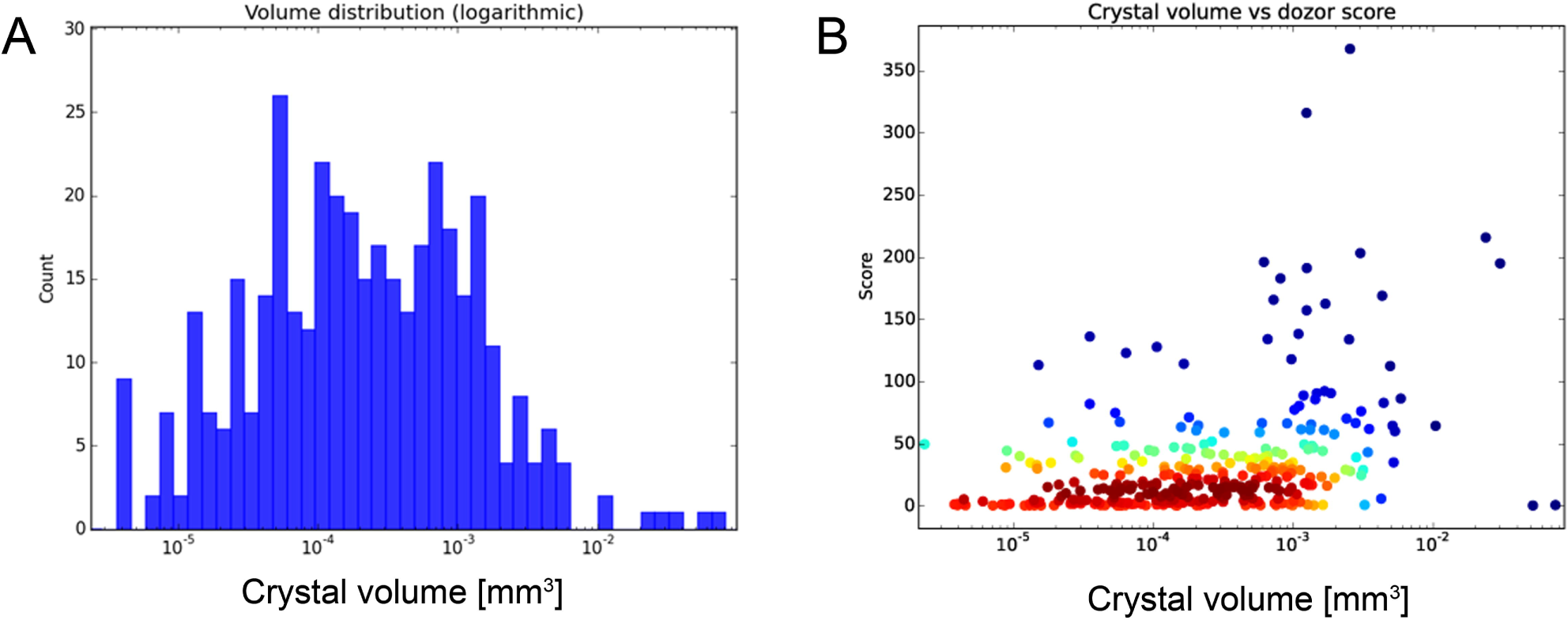
Volume distribution of crystals of afamin. A Distribution of volumes observed for afamin crystals and B the volume against the Dozor score (a measure of quality based on the radial intensity over background noise). The plot demonstrates the small number of crystals obtained that diffract to high resolution and the range of volumes obtained, requiring large scale screening.

It rapidly became apparent that widely varying cell parameters, leading to a variation in cell volumes of as much as 30-35%, were present in a primitive orthorhombic crystal form (*oP* lattice), in addition to several instances of a related primitive monoclinic cell (*mP*). While we were able to refine the monoclinic form (space group (SG) No. 4, *P*2_1_) with two molecules in the asymmetric cell as detailed earlier (Naschberger *et al.*, 2017), the merging statistics for the orthorhombic crystal forms (largely SG No. 18, *P*2_1_2_1_2) were generally poor. Despite nominal resolutions of around 2 Å or better the refinement stalled at high R-values without obvious avenues for improvement of the orthorhombic structure model. Dehydration experiments were therefore conducted on about one hundred orthorhombic crystals to determine if this form could be stabilised.

Afamin-gadoteridol complex crystals were screened on MASSIF-1 using automatic protocols for the location and optimal centring of crystals (Svensson *et al.*, 2015). The beam diameter was selected automatically to match the crystal volume of highest homogeneous quality (Svensson *et al.*, 2018). Strategy calculations accounted for flux and crystal volume in the parameter prediction for complete data sets. Despite appearing from different precipitant conditions, screening of 164 poorly diffracting crystals yielded highly anisotropic *oP* diffraction data similar to the previously obtained orthorhombic crystal form but with a significantly smaller unit cell. After pre-processing with XDSproc the data were submitted to the StarAniso server (Tickle *et al.*, 2018) and the R-free reflections from dehydrated orthorhombic afamin 6FAK were used for all orthorhombic data sets. Despite very poor merging statistics and low completeness (Table 1) the anisotropy-corrected data gave clear molecular replacement solutions in SG No. 18, *P*2_1_2_1_2, but with template 5OKL_A from monoclinic afamin (Naschberger *et al.*, 2017) using Phaser (McCoy *et al.*, 2007). Although no significant anomalous signal could be extracted from the poor data recorded at 12.835 keV (above the Gd L3-edge of 7.243 keV), strong positive difference peaks indicated the likely presence of 3 Gd sites, two of them in a pair exhibiting the same Gd-Gd distance (Figure 7) observed in high resolution models of lysozyme complexed with Gd-DO3A (Gorel *et al.*, 2017, Holton *et al.*, 2014, Girard *et al.*, 2002). Partial model building with Coot (Emsley *et al.*, 2010) and refinement with Refmac (Murshudov *et al.*, 2011) proceeded as described for the monoclinic form of afamin (Naschberger *et al.*, 2017). As a consequence of high data anisotropy, poor merging statistics, and low completeness, various disordered regions could not be modelled due to streaky and discontinuous maps. While core regions of afamin were well defined and had good geometry, R-free values never dropped below 34% and only an incomplete model could be obtained.

**Table 1.**
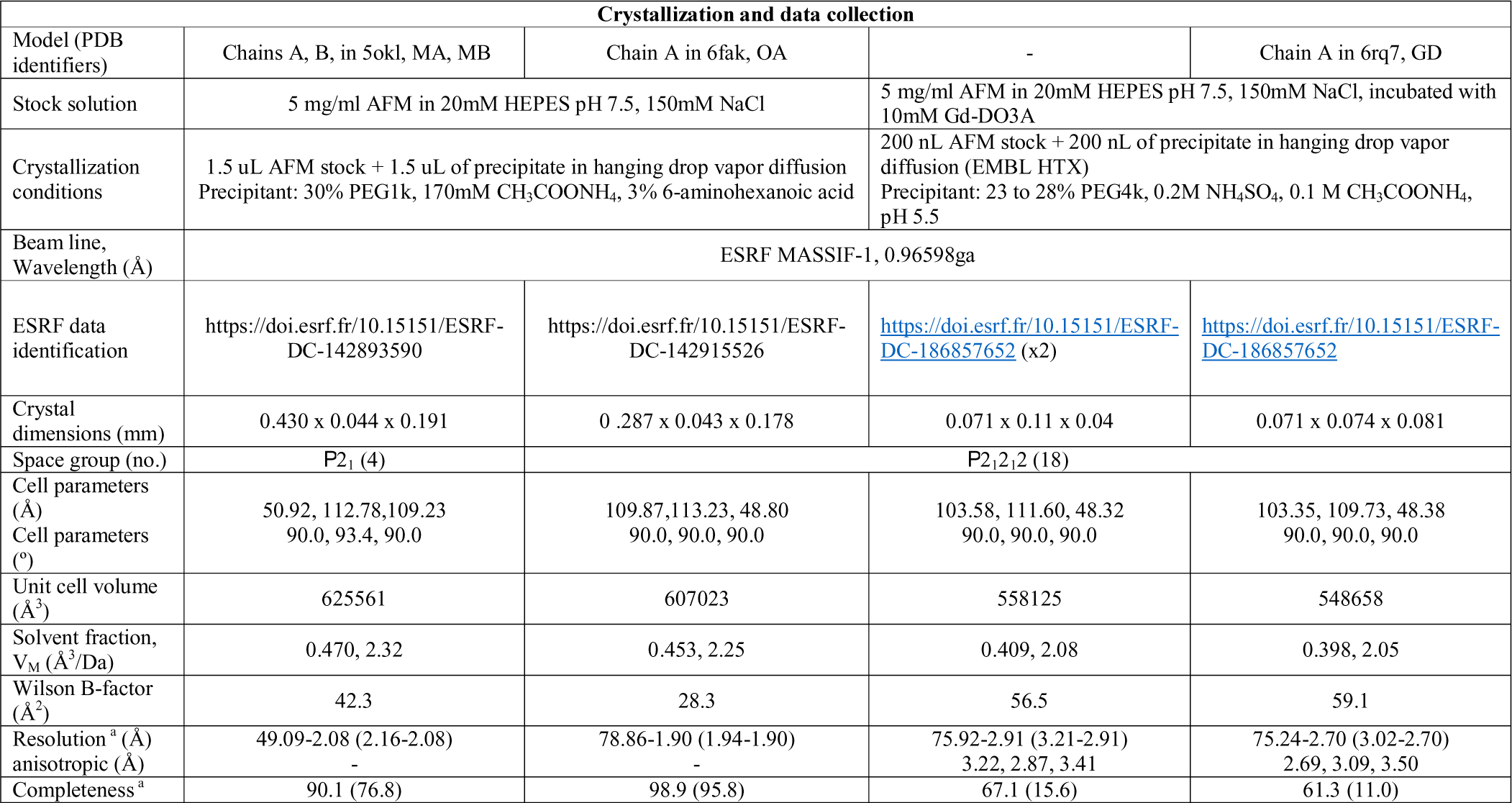
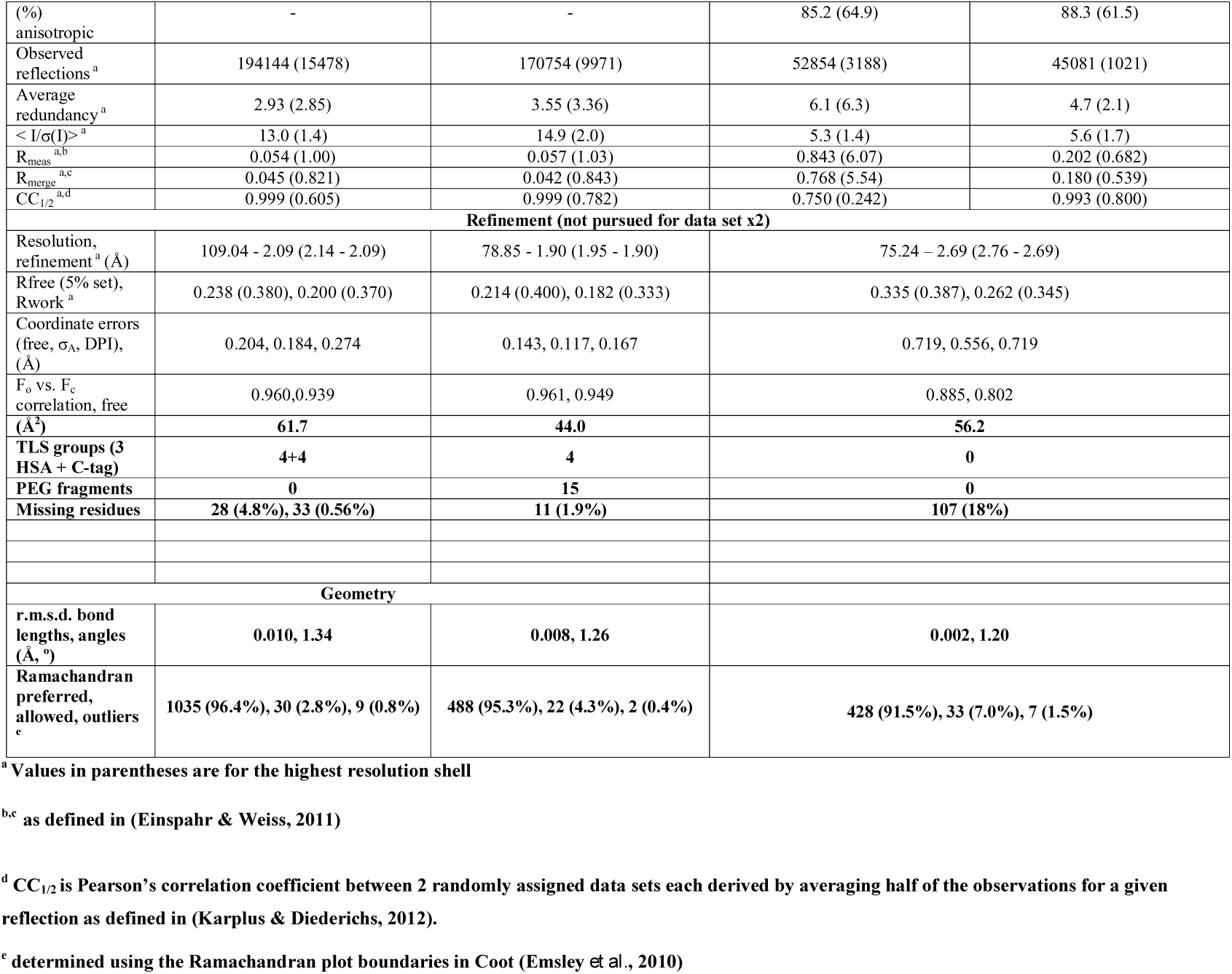
Crystallization, data collection, and refinement statistics for afamin structure models.

### 2.4 Dehydration experiments

Dehydration experiments were performed using an HC-Lab instrument (Arinax, France) (Sanchez-Weatherby *et al.*, 2009). The relative humidity (RH) in equilibrium with the mother liquor was predicted to be 98% (Bowler *et al.*, 2017, Wheeler *et al.*, 2012) and single crystals were mounted at this RH on micromesh mounts (Mitegen, Ithica, USA) on the RoboDiff goniometer (Nurizzo *et al.*, 2016) at the ESRF beamline MASSIF-1 (Bowler, Nurizzo, *et al.*, 2015). An automated workflow was then launched (Svensson *et al.*, 2015) that decreased the RH in steps of 1% with an equilibration time of 5 minutes with data collection and analysis at the end of each step. When the RH reached 85% a significant increase in diffraction quality was observed (Figure 3A and supplementary movie 1) accompanied by a reduction in the *a* cell edge and mosaic spread (Figure 3B). The resolution increase was maintained until a RH of 75% was reached, thereafter diffraction was lost (supplementary movie 1). This transition was confirmed by repeating the protocol on 5 crystals in the same manner. In order to optimise this transition, a number of protocols were attempted that also incorporated automated data collection procedures (Svensson *et al.*, 2015). A major bottleneck in refining the best protocol was the need to screen crystals before starting dehydration, therefore, in order to cover a wide range of conditions a multi-crystal protocol was used. Multiple *oP*-form crystals (3 to 7) were mounted on mesh loops (Figure 3B) and subjected to dehydration gradients to 80, 75 and 70% RH in either multiple or single steps, also varying final equilibration times (5 to 20 minutes). Once the protocols were completed, crystals were then cryocooled directly (Pellegrini *et al.*, 2011) and launched for autonomous characterisation and data collection using software and procedures (Figure 4B and C) described in §2.3 for multiple crystals (Svensson *et al.*, 2018). The obtained data sets were then analysed for quality and this information was fed back into subsequent rounds of dehydration with a refined protocol (Figure 3D). The final protocol used was to dehydrate crystals to an RH of 75% in a single step with a final equilibration time of 15 minutes. Three rounds of optimisation were performed using a total of 134 crystals. Using this procedure, the orthorhombic crystal form could be stabilised and data were collected that extended to a Bragg spacing of ~1.8 Å. The best data were manually reprocessed with the software described in §2.3 with statistics provided in Table 1. No twinning or pseudosymmetry (translational NCS) were detected in the data sets used for the dehydrated model refinement, and modest anisotropy in the orthorhombic data was accounted for in anisotropic scaling (Murshudov *et al.*, 2011) during refinement.

**Figure 3.**
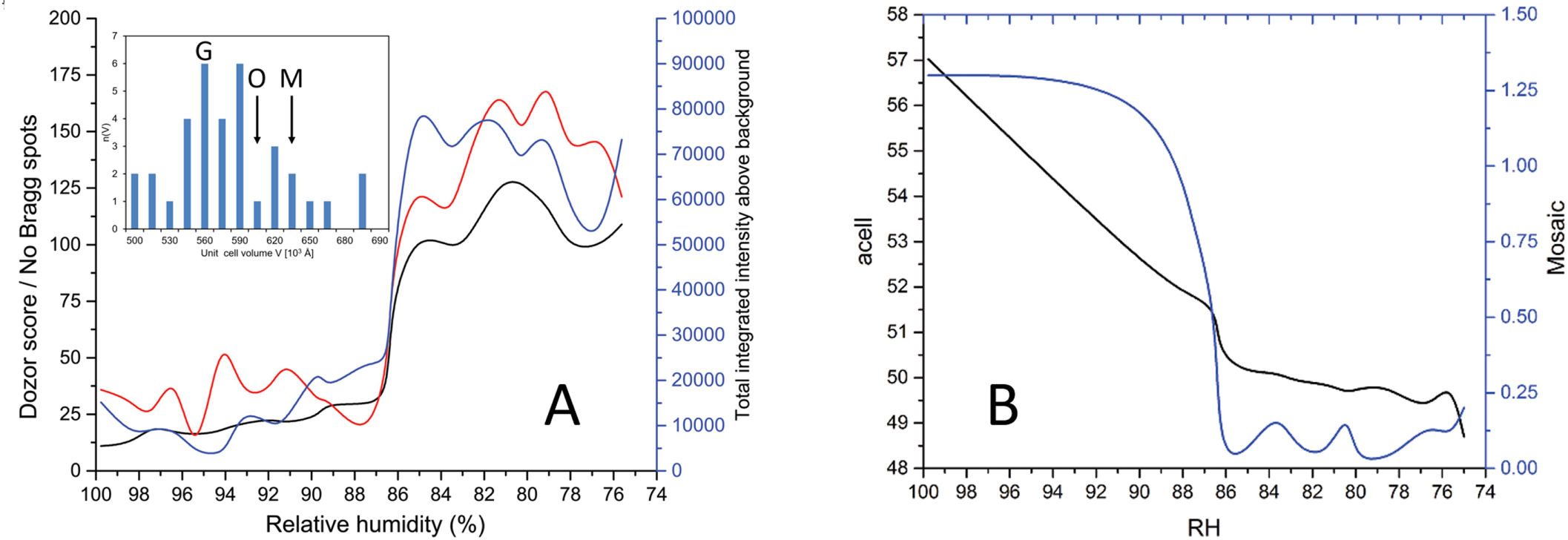
Effect of dehydration on the properties of afamin crystals. (A) Diffraction quality indicators increase dramatically around 86% relative humidity (RH). The insert shows the cell volumes of AFM crystal forms (M, monoclinic; O, orthorhombic dehydrated; G, Gd-DO3A-AFM complex). (B) The decrease in *a* cell dimension and mosaic spread on decreasing RH.

**Figure 4.**
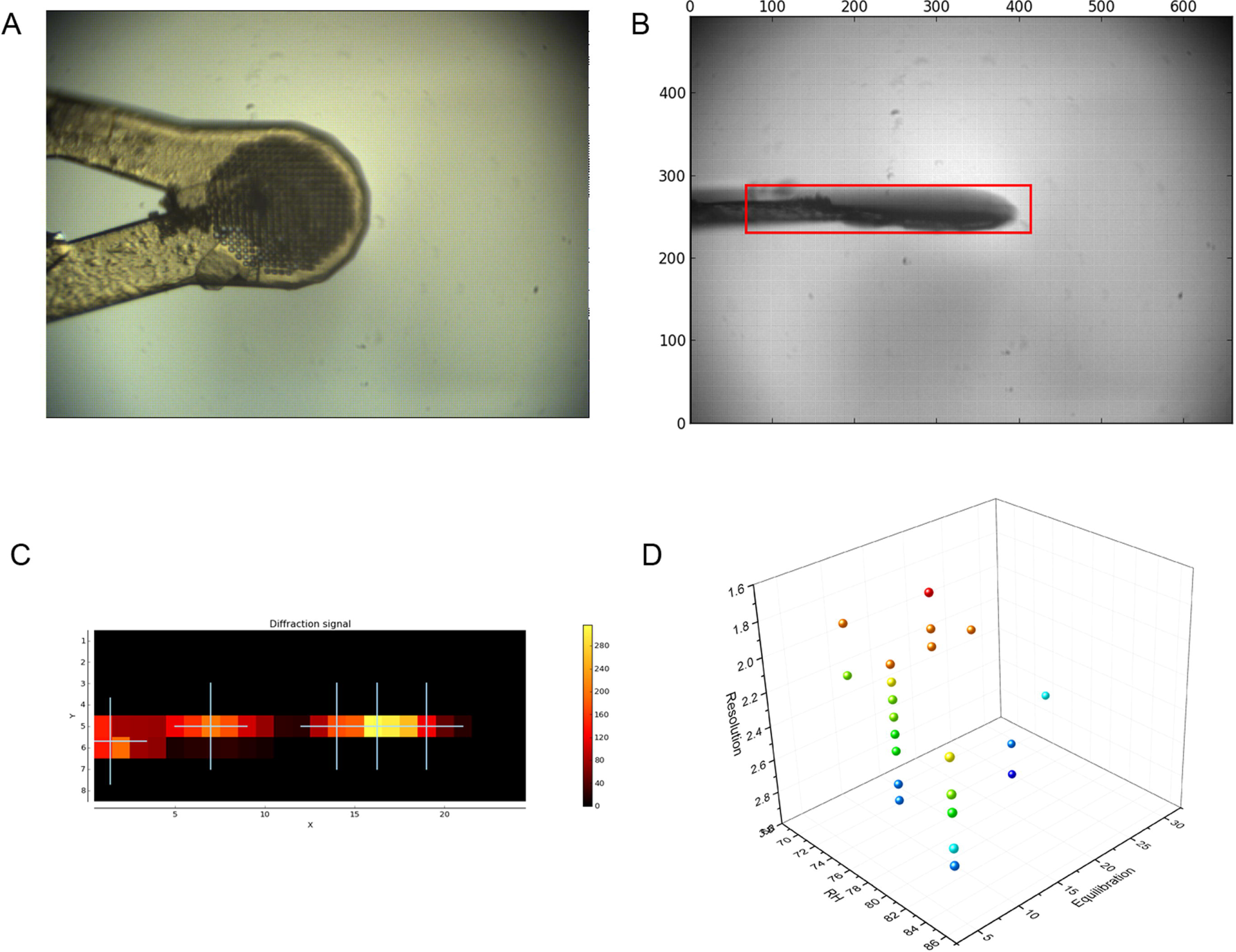
Multi-crystal strategy for optimising dehydration protocol. (A) Multiple crystals were loaded onto micro-mesh mounts and then subjected to various dehydration protocols and cryo-cooled directly. (B and C) Multi-crystal data collection protocols were then run on the supports on MASSIF-1 allowing the protocols to be assessed despite crystal variation. (D) Finally, the results could be assessed showing which protocol should be used and applied to further crystals.

## 3.0 Results and discussion

### 3.1 Effect of dehydration on crystal packing

The change between the monoclinic and orthorhombic cells is subtle. The 2-fold NCS axis in the monoclinic form is very close to the crystallographic 2-fold axis in the orthorhombic form, illustrated by comparing the self-rotation function between the two forms, and appearance of a NCS peak in the native Patterson map (Figure 5). The symmetry gain towards the orthorhombic form may be energetically favourable but many of the crystals tested may not have completed the transition completely, leading to heterogeneity. This may be due to slow or incomplete dehydration in the native crystallization drops or due to packing defects as the crystals grow. The use of a humidity control device has allowed the orthorhombic crystal form to be stabilised. The transformation involves a contraction in the *a* cell edge of over 10 Å and the β angle by ~3° to 90° (Figure 6 and supplementary movie 2). This change shifts one molecule in the asymmetric unit allowing it to become related by crystallographic symmetry to its mate, reducing the asymmetric unit to a single molecule in the orthorhombic cell (Figure 6). This movement also stabilises certain loop regions that were not visible and could not be modelled in the monoclinic structures. The number of crystal contacts between symmetry related molecules increases from 1999 to 2752, an increase of 37%. The result is a significant stabilisation of the molecule that can be seen in the B-factors (Figure 6). This transition from *mP* to *oP* lattice takes place in the same crystal, in contrast to the formation of the smaller orthorhombic AFM-Gd-DO3A complex crystals grown with slightly different precipitates and at different pH (6.5 vs. 4.6).

**Figure 5.**
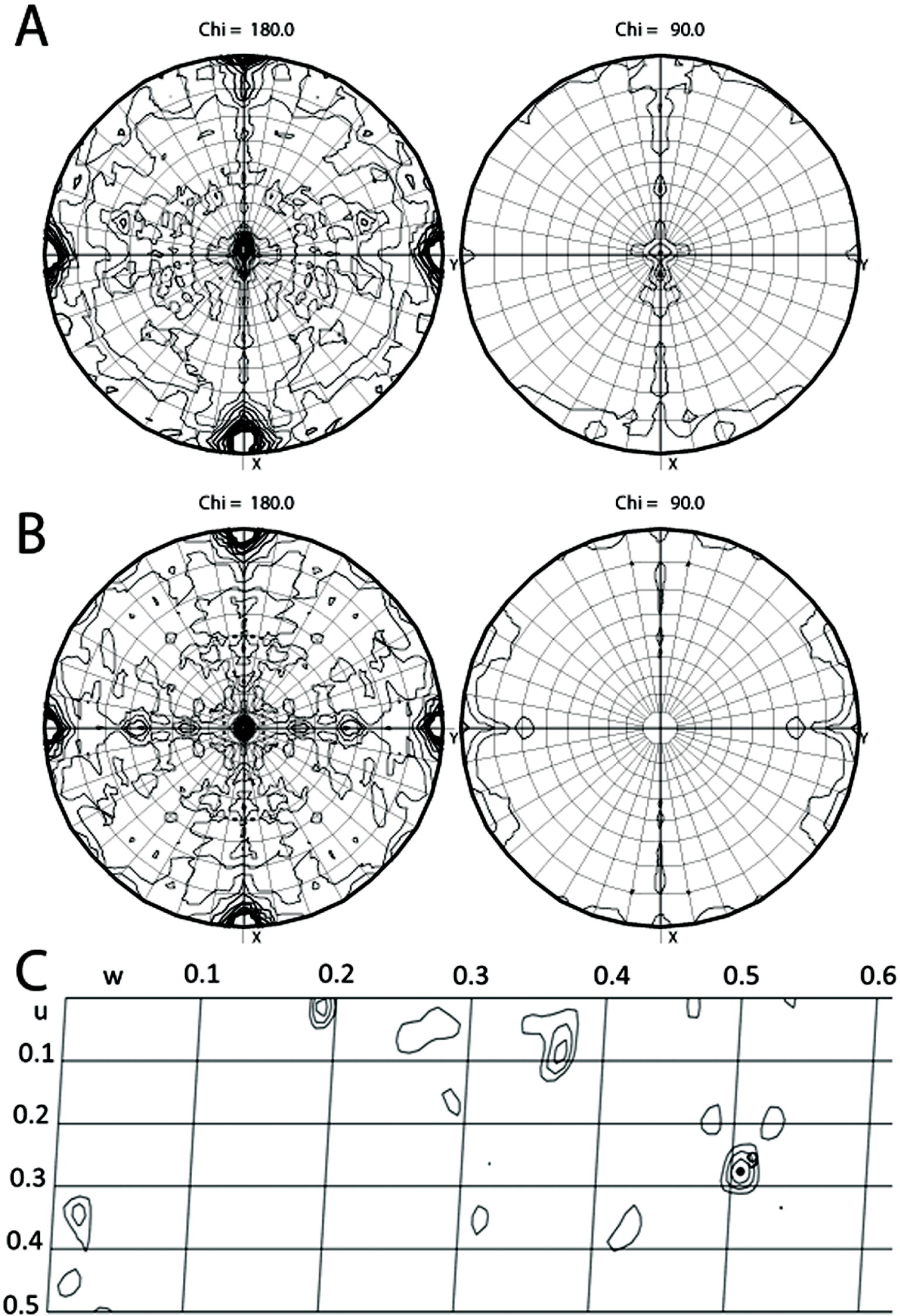
Self-rotation functions and a native Patterson map of the monoclinic and orthorhombic crystal forms show the close relation between NCS and CS. The self-rotation function shows the peaks from the non-crystallographic symmetry in the monoclinic form (A) are very close to those for the crystallographic symmetry in the orthorhombic crystal (B) implying only a small shift is needed to satisfy the requirements for higher symmetry. While the orthorhombic map (B) displays almost perfect mm symmetry, reduced symmetry along Y in (A) becomes visible. The Harker section (u,1/2, w) of the native Patterson map (C) shows a weak peak at u,v,w = (0.277, ½, 0.507) indicating the location of the 3.7° tilted NCS axis originating from true crystallographic axis at u,v w = (¼, ½, ½).

**Figure 6.**
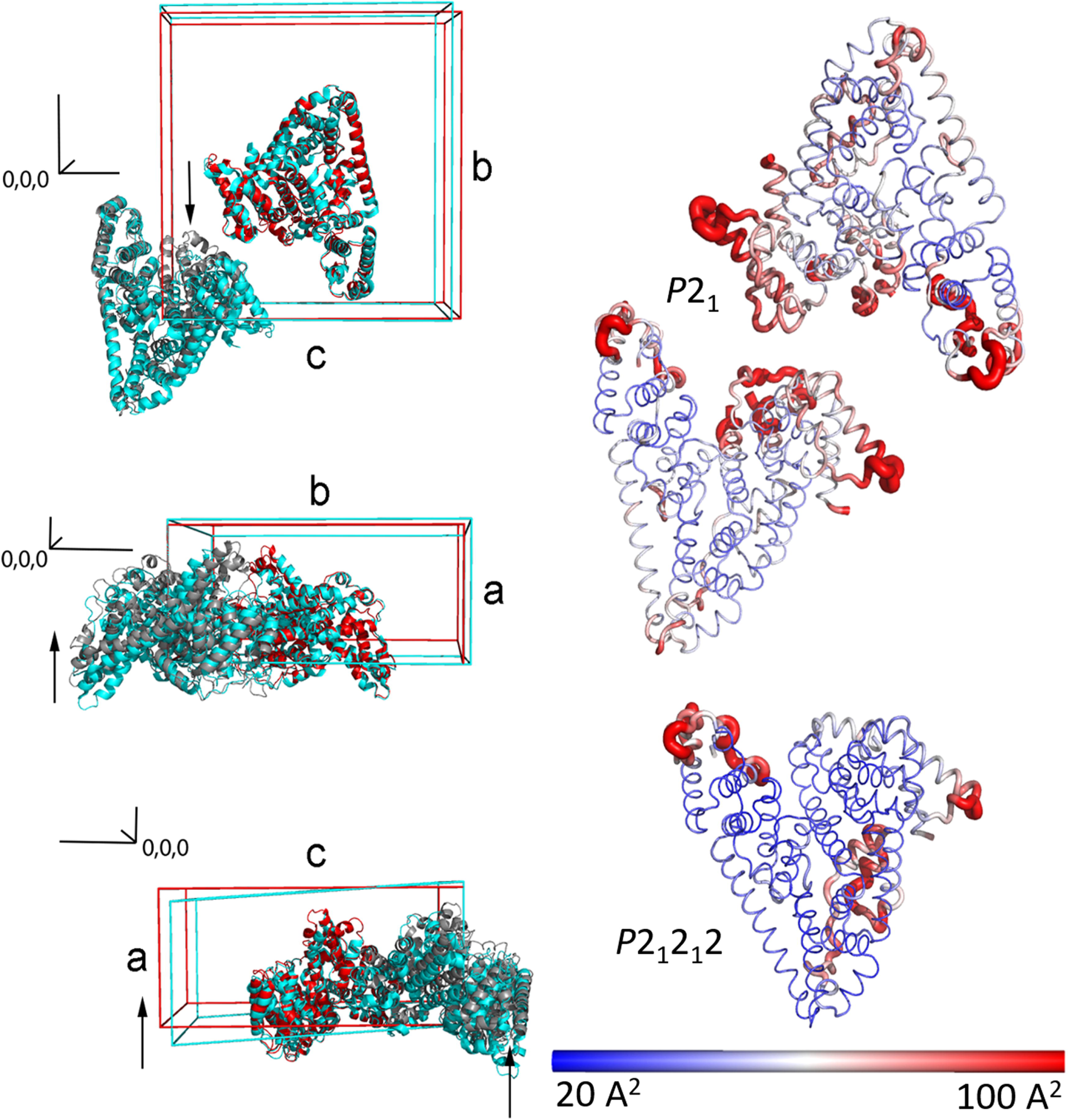
Packing and flexibility changes induced by dehydration. Left column: Dehydration of the monoclinic cell (cyan) leads to a decrease in unit cell volume and a reduction of the β angle to 90°, the latter has the most profound effect moving one molecule in the asymmetric unit by up to 10 Å. This movement reduces the asymmetric unit to a single molecule in an orthorhombic form (red, symmetry related mate in grey) and results in the stabilisation of several loop regions where electron density was not visible previously. Note: the axis labels refer to the monoclinic cell. In order to demonstrate the shift in unit cells the orthorhombic form was reindexed to match the monoclinic convention. Therefore, for the orthorhombic cell the conversion is *a → c* and *c → a*. Right column: Cartoon representation of the asymmetric units of the monoclinic versus orthorhombic crystal forms. The Cα chains are shown coloured by B-factor indicating a reduction in flexibility in multiple regions.

The cell volume distribution of data sets that could be indexed (0A insert) shows that as expected under dehydration the volume shrinks about 3.5%. An even more dramatic cell contraction dominated by a further 6Å contraction in *a* compared to the isomorphous orthorhombic dehydrated crystals and an almost 13% decrease form the monoclinic cell volume occurs in the AFM-GdDO3A complex crystals. Whether this decrease in cell volume is mediated by the single Gd-DO3A molecule located at a crystal contact (Figure 7) or a result of shifting local charge distributions due to the lower crystallization pH of 5.5 vs 6.5 for the native AFM remains unknown.

**Figure 7.**
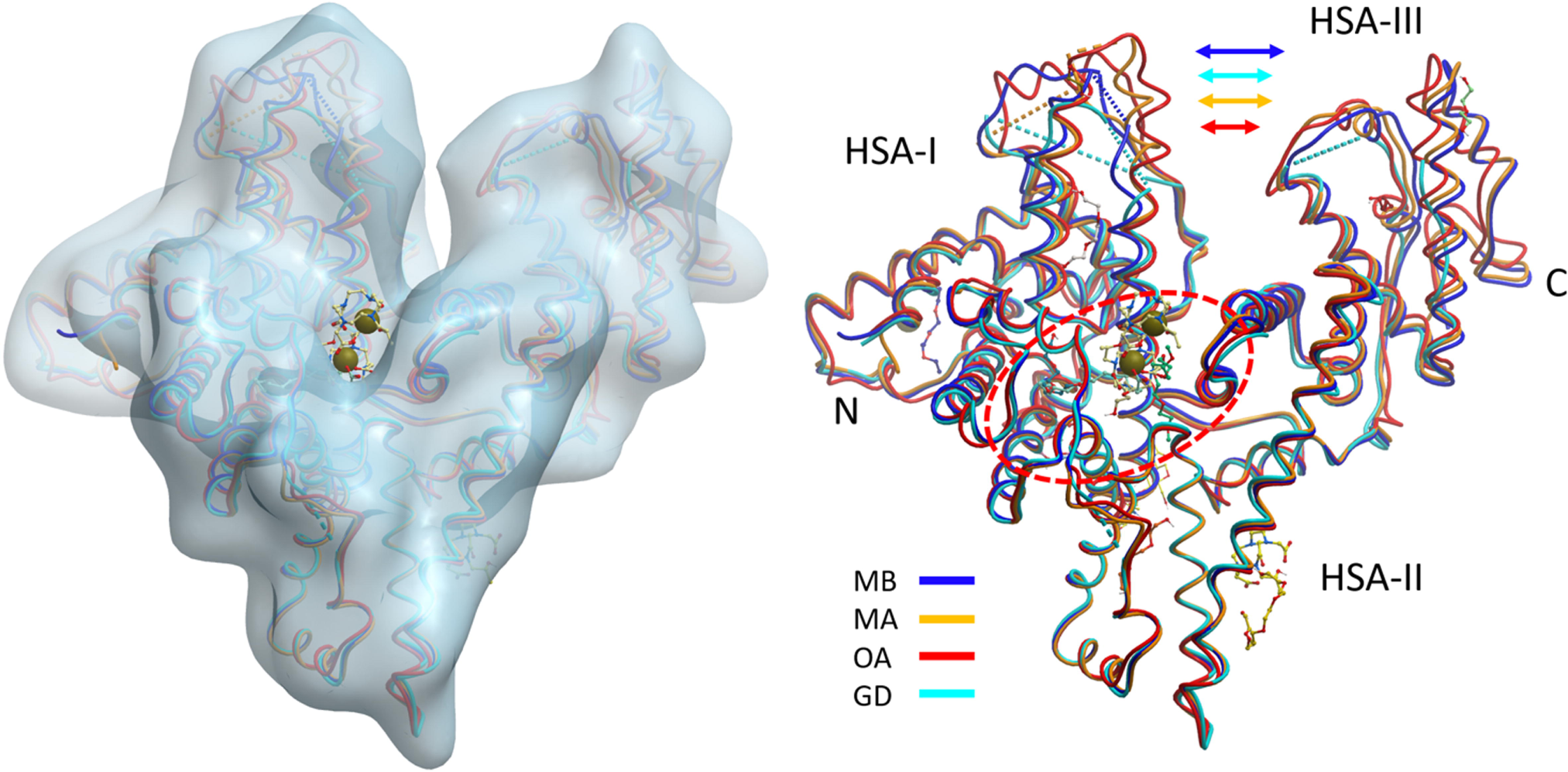
Overview of the afamin structure models. (A) The major binding site of afamin, corresponding to the Sudlow S1 drug binding site in albumin, is located in the center of the heart-shaped molecule between the equivalent of HSA domains I and III. The entrance of the deep hydrophobic cleft harbours the Gd-DO3A chelate molecules indicated by the grey spheres. (B) The four structure models of afamin (monoclinic model MA, 5okl_A; monoclinic model MB, 5okl_B; the dehydrated orthorhombic model OA, 6fak_A; and the orthorhombic form of the Gd-DO3A complex, GD) are superimposed on the HSA-II domain. The motion of the domains I and III towards each other affects access to the binding site. Shown in ball and stick model are also the first hexose molecules of the paucimannose basic glycosylations and PEG molecules of dehydrated model 6fak_A. Figure produced with Molsoft ICMPro (Abagyan *et al.*, 2006).

### 3.2 Contextual flexibility affects binding site analysis

One of the most interesting insights gained from the comparison of four different AFM structure models (monoclinic model MA, 5okl_A; monoclinic model MB, 5okl_B; the dehydrated orthorhombic model OA, 6fak_A; and the orthorhombic Gd-DO3A complex, GD; cf. section 3.4) is that the relative motions of the domains forming the deep cleft in the centre of the heart shaped molecule (following the HSA description) do affect the shape of the deep hydrophobic binding pocket at the equivalent of the HSA Sudlow 1 (S1) drug binding site (Naschberger *et al.*, 2017). The breathing motions that occur between a transition from the more open MB conformation to the tighter MA and OA conformations (Figure 7) also affect the S1 binding site. In the dehydrated OA form an additional minor pocket opens up extending the primary S1 binding cleft, which is then occupied with what we believe is a PEG fragment (cf. also section 3.3, Figure 8).

**Figure 8.**
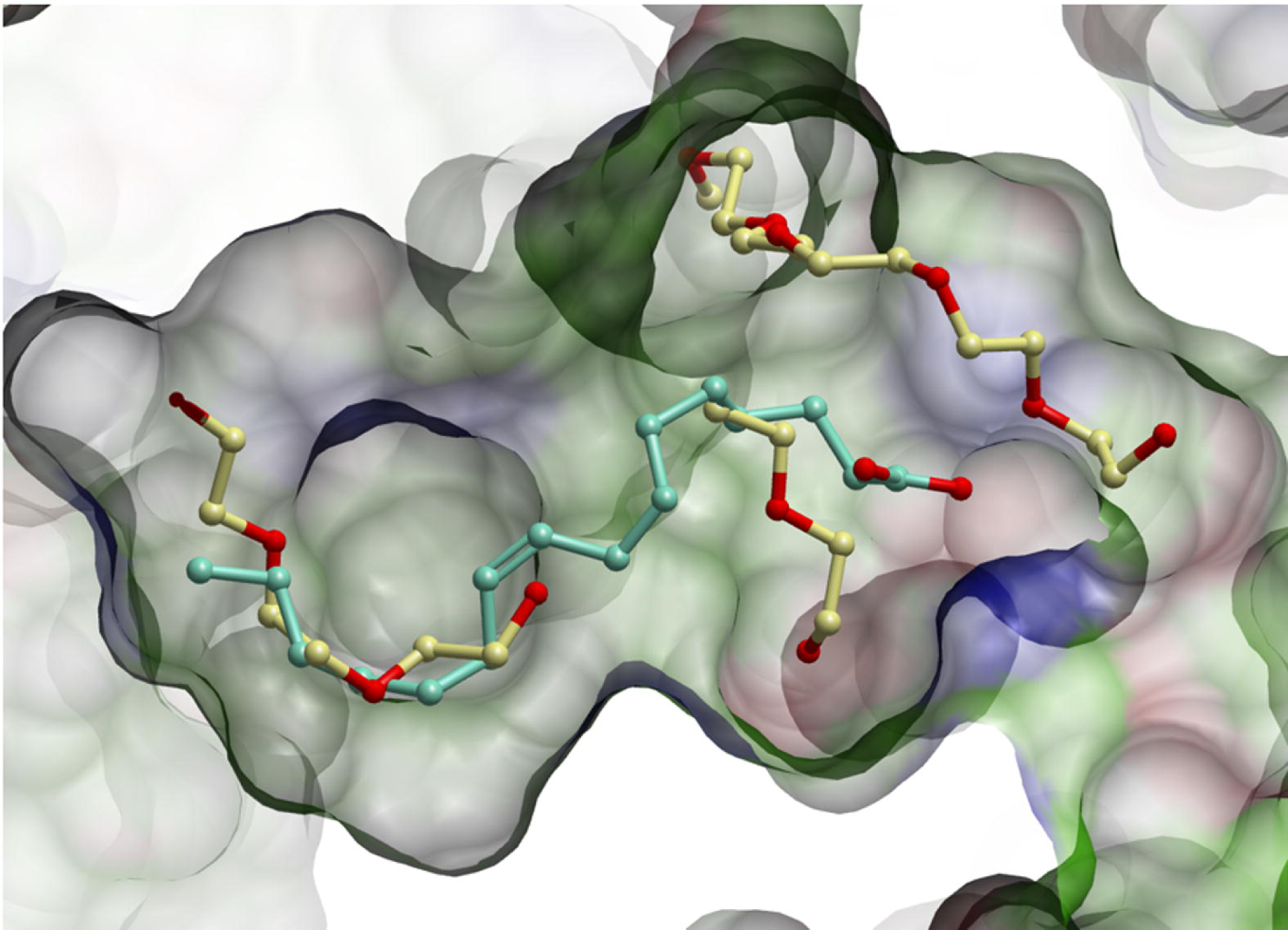
Detail view of the hydrophobic binding cleft in the dehydrated crystal structure of afamin. Shown is a cross section of the deep hydrophobic binding pocket (left side) over a transparent property-coloured residue surfaces (white: lipophilic; green aromatic lipophilic; red: hydrogen bonding acceptor potential; blue: hydrogen bond donor potential). Overlaid is in green ball and stick model the palmitoleic acid as modelled into the cavity of 5okl_B (MB). Shown in yellow ball and stick presentation are PEG fragments modelled into density of dehydrated structure 6fak (OA). Visible is a narrow second channel (top middle of the figure) harbouring a long PEG fragment in 6fak. Figure produced with Molsoft ICMPro (Abagyan *et al.*, 2006).

While one can argue that the OA binding site of the dehydrated crystal structure does not represent a native solution conformation, it does amplify the concerns that a single crystal structure, particularly in a restrictive crystal packing, may not be sufficient as a basis for a drug lead discovery study (Dym *et al.*, 2016). The power of multiple crystals forms to explore the conformational space of a protein in its native solvent environment has repeatedly been made, e.g. (Naschberger *et al.*, 2016). Given modern high throughput crystallography methods, as many different crystal forms should be examined as possible in order to obtain a complete picture, particularly in the case of highly promiscuous small molecule and drug transporters as exemplified by albumin and afamin. The frequent presence of PEG molecules in crystallization cocktails adds an additional level of uncertainty to structure guided drug lead discovery (Dym *et al.*, 2016).

### 3.3 Probing the binding site for hydrophobic molecules: different crystal forms can lead to different ligand occupancies

The most prominent feature of the AFM molecule is the deep, central binding cavity extending almost across the entire molecule (Figure 8). While the solvent exposed region of this deep cavity (where the Gd-DO3A is located) is in the vicinity of the HSA Sudlow 1 drug binding site, the inner lining of the in AFM much deeper pocket is almost exclusively formed by hydrophobic and lipophilic residues. This deep hydrophobic pocket is likely the key anchor for the interaction of palmitoylated Wnt3a with afamin (Mihara *et al.*, 2016), and distinct electron density in this cavity of the non-dehydrated monoclinic model chain B was attributed to palmitoleic acid (PAM) and supported by lipid analysis of the purified and crystallized AFM sample (Naschberger *et al.*, 2017). Additional supporting evidence for fatty acid binding such as lipid analysis and a careful analysis of the chemical site environment is almost always necessary when PEGs are a component of the crystallization cocktail, because at common resolutions (around 2.5-2.0 Å) electron density shape alone does not allow clear distinction between an aliphatic fatty acid tail and a PEG molecule.

While the frequent presence of high concentrations of PEG molecules in crystallization cocktails adds uncertainty to the identification of ligands - in particular in the case of fatty acid chains - the possibility of forcing by dehydration PEG molecules into binding sites as surrogate probes is intriguing. In the example of afamin, the presence of the secondary binding channel occupied with a PEG molecule (Figure 8) opens up the speculation that afamin could – at least from a structural point of view - also accommodate moieties with two fatty acid chains such as phosphatidylcholines.

### 3.4 Gd-DO3A in afamin: The case for salvaging poor data and how to disseminate them

Following conventional wisdom, the data collection statistics in Table 1 (x2 and x3) would suggest that both tabulated Gd data sets are utterly useless. By historical standards, the merging statistics are horrendous, the data are highly anisotropic, and the completeness unacceptably low. As expected from these statistics, we were unable to extract any anomalous difference signal, which is expected to be a respectable 3.4% at 12.835 keV (0.9660Å) and thus detectable with reasonable data quality (Lemke *et al.*, 2002) and full occupancy given the Gd L3-edge of 7.243 keV. In absence of anomalous signal and in view of the abysmal quality metrics, the data would have been discarded by default. Nonetheless, we were able to extract useful and informative information from the data.

Contrary to expectations, molecular replacement with Phaser (McCoy *et al.*, 2007) using data corrected for anisotropy with StarAniso (Tickle *et al.*, 2018) yielded clear and unique solutions. A best solution was obtained with monoclinic AFM search model MA (5okl_A, TFZ 27.9, R-free 0.48), while the worst solution was obtained with the dehydrated orthorhombic AFM model OA (6fak_A, TFZ 22.5, R-free 0.52). The reason might be that the domain arrangements of HAS-I and HAS-III (Figure 7) in MA and the Gd model are similar. After MR and a first round of restrained refinement three large positive difference electron density peaks became immediately prominent (Figure 9) in maps based on both data sets. Smaller peaks varied between the two maps and were mostly located in unmodelled or incorrectly modelled parts of the map.

**Figure 9.**
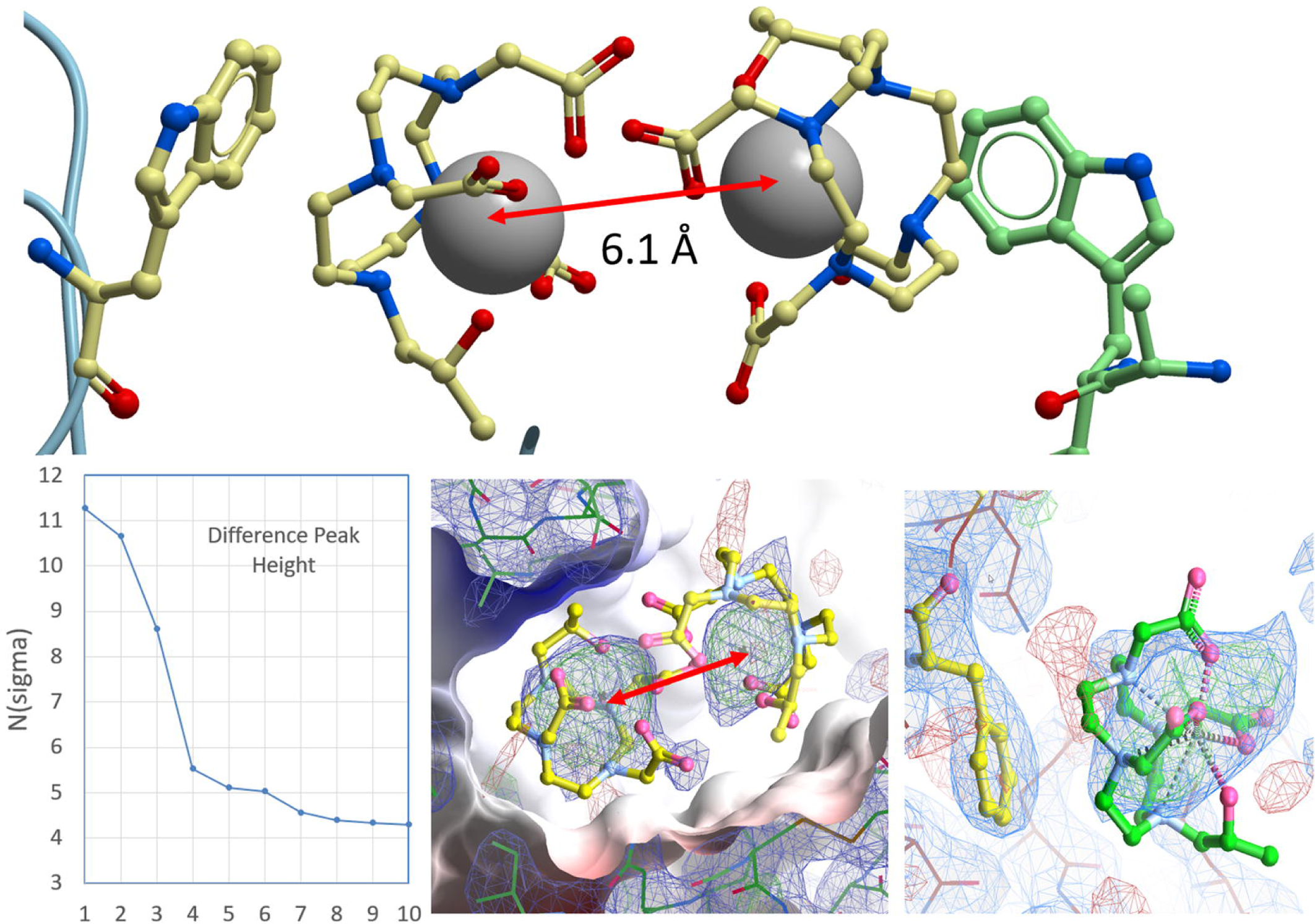
Gd-DO3A sites in lysozyme and afamin. The top panel shows the arrangement of the Gd-DO3A sites in a high-resolution model 1h87 of lysozyme (Girard *et al.*, 2003) where the Gd pair forms a crystal contact involving hydrophobic interaction of the DO3A methyl groups with W62 of one molecule and W123 of a symmetry mate. A similar arrangement exists in afamin, where two of the strongest difference peaks (insert) appear in the central hydrophobic binding cleft. Placed Gd atoms refined with an occupancy between 0.9 and 0.7, and the distance between them is also 6.1 Å as in the lysozyme model, with enough space in the binding site to accommodate the DO3A crowns of the Gd-DO3A complexes. The bottom centre panel shows m*F*_o_-D*F*_c_ positive omit difference electron density displayed at a 2.5 σ level in green and 2m*F*_o_-D*F*_c_ density displayed at 0.8 σ in blue. The DO3A crowns are not refined but placed in an orientation corresponding to 1h87. In afamin, the methyl groups of the modelled DO3A crown face towards a phenylalanine lining the bottom (left side of the figure) of the binding pocket, resembling the interactions observed in 1h87. The same interaction can be proposed for the third Gd site located at a crystal contact in Afamin (bottom right). Top figure produced with Molsoft ICMPro (Abagyan *et al.*, 2006) and electron density figures produced with Coot (Emsley *et al.*, 2010).

After a few rounds of rebuilding it became clear that large parts of the model were disordered. Due to streaking and discontinuity in many parts of the map the model could not be completed, and in agreement with the data statistics, maps from data x3 appeared subjectively better than maps from x2. Refinement of X3 stalled at R-free values of ~ 0.36 despite good geometry for the traceable part of the model. At this point, Gd atoms were placed with occupancy set to 0.7 so that B-factors refined to values of about 150% of the mean environment B-factor. Subsequent occupancy refinement indicated Gd occupancies between 0.9 and 0.7. The protrusions and difference density in the large density blobs suggested additional features around the Gd, but we were unable to determine and refine the orientation of the DO3A ‘crown’ around the Gd-DO3A. However, the refined distance of 6.1 Å between the two Gd peaks corresponds exactly to the distance observed in the high resolution lysozyme structures, and we placed – but did not refine - the DO3A crowns in the two poses determined in 1h87 (Girard *et al.*, 2003). There were no collisions, and the proposed interaction of the 8 methyl groups of the DO3A crown with an aromatic residue lining the bottom of the binding pocket is similar to the one observed in lysozyme. The same holds for the third Gd-DO3A located at an AFM crystal contact (Figure 9).

### 4. A case for depositing anisotropic models – but how?

Our proposition of Gd-DO3A being present in AFM is based on preponderance of evidence in the form of plausibility arguments, augmenting the reasonable evidence of strong difference density. This is certainly far from proof beyond reasonable doubt (which anomalous methods would have provided) but plausible enough to suggest further studies particularly regarding the structural basis of transport of contrast agents by albumin (Caravan, 2009). Despite the poor data processing and refinement statistics we decided to submit the incomplete and anisotropic model to the PDB. The original diffraction images for all structures described here, in line with recent IUCr policy developments (Helliwell *et al.*, 2019), are also available for download from the ESRF data portal (Naschberger *et al.*, 2018a and 2018b; Naschberger *et al.*, 2019).

While our modest claim of Gd being present is reasonably plausible and of general interest, a valid question is whether the deposition of an incomplete, and lacking in details of the binding site, model would perhaps contaminate the PDB (and likely invite the ire of rote statistics data-miners, a problem we have cautioned against repeatedly (Wlodawer *et al.*, 2018, Weichenberger *et al.*, 2017). This concern largely originates from the fact that descriptors for highly anisotropic data and models cannot adequately deposited with the PDB, and thus lead to misleading statistics. With the increasing success of anisotropy correction methods and corresponding servers (Strong *et al.*, 2006, Tickle *et al.*, 2016) the limitations of isotropic, scalar presentation of diffraction data metrics are inadequate and outdated (Rupp, 2018). Our Gd model deposition illustrates the case.

The validation report for the Gd model, states an automatically extracted resolution limit of 2.69 Å. This is misleading given the anisotropy and the correspondingly low completeness (11% spherical in the highest resolution bin, Table 1) of the data. User expectations of model quality based on an isotropic model resolution of 2.69 Å are cannot be met, as would extrapolations of its usability for other purposes than the support for Gd presence. At minimum, there needs to be a means of depositing – or preferably, extracting from the respective anisotropy server logs or unmerged original diffraction data - elliptic resolution and completeness limits. Suggestions for data presentation and the annotation of consequences for the overall limitations of model anisotropy have been made (Rupp, 2018). The anisotropy reported in section 4 of the report likely gets overlooked and is also based on the submission of already elliptically corrected data and thus not sufficient.

What is noteworthy is that in contrast to absurdly high merging R-values precluding any acceptance of a structure model based on the Table 1 statistics, the CC_1/2_ seems to be a robust measure indicating that such data in principle can be used to extract at least some useful and valid information. Finally, it would be useful to allow annotation by the depositor of unusual features, restrictions, or warnings of shortcoming at the time of model deposition. A clear reference to such user-defined limitations, akin to the caveat statements, should be a prominent and data minable element in the final deposition.

## Acknowledgements

Work supported by the Austrian Science Fund (FWF) under project P28395-B26 to BR. We thank the HTX laboratory (EMBL Grenoble) for tireless assistance with crystallisation experiments and harvesting with the CrystalDirect robot.

## References

Abagyan, R., Lee, W. H., Raush, E., Budagyan, L., Totrov, M., Sundstrom, M. & Marsden, B. D. (2006). TIBS, 31, 76–78.

Adams, P. D., Afonine, P. V., Bunkóczi, G., Chen, V. B., Echols, N., Headd, J. J., Hung, L.-W., Jain, S., Kapral, G. J., Grosse Kunstleve, R. W., McCoy, A. J., Moriarty, N. W., Oeffner, R. D., Read, R. J., Richardson, D. C., Richardson, J. S., Terwilliger, T. C. & Zwart, P. H. (2011). Methods 55, 94–106.

Altamirano, A., Naschberger, A., Fürnrohr, B. G., Saldova, R., Struwe, W. B., Jennings, P. M., Millán Martín, S., Malic, S., Plangger, I., Lechner, S., Pisano, R., Peretti, N., Linke, B., Aguiar, M. M., Fresser, F., Ritsch, A., Lenac Rovis, T., Goode, C., Rudd, P. M., Scheffzek, K., Rupp, B. & Dieplinger, H. (2018). J. Proteo. Res. 17, 1269–1277.

Bowler, M. G., Bowler, D. R. & Bowler, M. W. (2017). J. Appl. Cryst. 50, 631–638.

Bowler, M. W., Montgomery, M. G., Leslie, A. G. & Walker, J. E. (2006). Acta Cryst. D62, 991–995.

Bowler, M. W., Mueller, U., Weiss, M., Sanchez-Weatherby, J., Sorensen, T., Thunnissen, M., Ursby, T., Gobbo, A., Russi, S., Bowler, M. G., Brockhauser, S., Svensson, O. & Cipriani, F. (2015). Cryst. Growth. Des. 15, 1043–1054

Bowler, M. W., Nurizzo, D., Barrett, R., Beteva, A., Bodin, M., Caserotto, H., Delageniere, S., Dobias, F., Flot, D., Giraud, T., Guichard, N., Guijarro, M., Lentini, M., Leonard, G. A., McSweeney, S., Oskarsson, M., Schmidt, W., Snigirev, A., von Stetten, D., Surr, J., Svensson, O., Theveneau, P. & Mueller-Dieckmann, C. (2015). J. Sync. Rad. 22, 1540–1547.

Caravan, P. (2009). Acc Chem Res 42, 851–862.

Cramer, P., Bushnell, D. A., Fu, J. H., Gnatt, A. L., Maier-Davis, B., Thompson, N. E., Burgess, R. R., Edwards, A. M., David, P. R. & Kornberg, R. D. (2000). Science 288, 640–649.

Dieplinger, H., Ankerst, D. P., Burges, A., Lenhard, M., Lingenhel, A., Fineder, L., Buchner, H. & Stieber, P. (2009). Cancer Epidemiology Biomarkers & Prevention 18, 1127–1133.

Dym, O., Song, W., Felder, C., Roth, E., Shnyrov, V., Ashani, Y., Xu, Y., Joosten, R. P., Weiner, L., Sussman, J. L. & Silman, I. (2016). Protein Sci. 25, 1096–1114.

Einspahr, H. M. & Weiss, M. S. (2011). International Tables for Crystallography F, 2nd Ed., 2nd Ed. ed., edited by E. Arnold, D. M. Himmel & M. G. Rossmann, pp. 64–74. Chichester: Wiley.

Emery, V. C. (1991). Practical Molecular Virology: Viral Vectors for Gene Expression, edited by M. K. L. Collins, pp. 287–307. Totowa, NJ: Humana Press.

Emsley, P., Lohkamp, B., Scott, W. G. & Cowtan, K. (2010). Acta Cryst. D66, 486–501.

Fasano, M., Curry, S., Terreno, E., Galliano, M., Fanali, G., Narciso, P., Notari, S. & Ascenzi, P. (2005). IUBMB Life 57, 787–796.

Girard, E., Chantalat, L., Vicat, J. & Kahn, R. (2002). Acta Cryst. D58, 1–9.

Girard, E., Stelter, M., Anelli, P. L., Vicat, J. & Kahn, R. (2003). Acta Cryst. D59, 118–126.

Gorel, A., Motomura, K., Fukuzawa, H., Doak, R. B., Grünbein, M. L., Hilpert, M., Inoue, I., Kloos, M., Kovácsová, G., Nango, E., Nass, K., Roome, C. M., Shoeman, R. L., Tanaka, R., Tono, K., Joti, Y., Yabashi, M., Iwata, S., Foucar, L., Ueda, K., Barends, T. R. M. & Schlichting, I. (2017). Nature commun. 8, 1170.

Helliwell, J. R., Minor, W., Weiss, M. S., Garman, E. F., Read, R. J., Newman, J., van Raaij, M. J., Hajdu, J. & Baker, E. N. (2019). Acta Cryst. D75, 455–457.

Heras, B. & Martin, J. L. (2005). Acta Cryst. D61, 1173–1180.

Holton, J. M., Classen, S., Frankel, K. A. & Tainer, J. A. (2014). FEBS J. 281, 4046–4060.

Hu, N. J., Iwata, S., Cameron, A. D. & Drew, D. (2011). Nature 478, 408–411.

Kabsch, W. (2010). Acta Cryst. D66, 125–132.

Kadlec, J., Hallacli, E., Lipp, M., Holz, H., Sanchez-Weatherby, J., Cusack, S. & Akhtar, A. (2011). Nature Struct. Mol. Biol. 18, 142–149.

Karplus, P. A. & Diederichs, K. (2012). Science 336, 1030–1033.

Kronenberg, F., Kollerits, B., Kiechl, S., Lamina, C., Kedenko, L., Meisinger, C., Willeit, J., Huth, C., Wietzorrek, G., Altmann, M. E., Thorand, B., Melmer, A., Dähnhardt, D., Santer, P., Rathmann, W., Paulweber, B., Koenig, W., Peters, A., Adham, I. M. & Dieplinger, H. (2014). Circulation: Cardio. Genet. 7, 822–829.

Lemke, C. T., Smith, G. D. & Howell, P. L. (2002). Acta Cryst. D58, 2096–2101.

McCoy, A. J., Grosse-Kunstleve, R. W., Adams, P. D., Winn, M. D., Storoni, L. C. & Read, R. J. (2007). J. Appl. Cryst. 40, 658–674.

Mihara, E., Hirai, H., Yamamoto, H., Tamura-Kawakami, K., Matano, M., Kikuchi, A., Sato, T. & Takagi, J. (2016). Elife 5, e11621.

Monaco, S., Gordon, E., Bowler, M. W., Delageniere, S., Guijarro, M., Spruce, D., Svensson, O., McSweeney, S. M., McCarthy, A. A., Leonard, G. & Nanao, M. H. (2013). J. Appl. Cryst. 46, 804–810.

Murshudov, G. N., Skubak, P., Lebedev, A. A., Pannu, N. S., Steiner, R. A., Nicholls, R. A., Winn, M. D., Long, F. & Vagin, A. A. (2011). Acta Cryst. D67, 355–367.

Naschberger, A., Fürnrohr, B., Lenac Rovis, T., Malic, S., Scheffzek, K., Dieplinger, H. & Rupp, B. (2016). Acta Cryst. D72, 1267–1280.

Naschberger, A., Orry, A., Lechner, S., Bowler, M. W., Nurizzo, D., Novokmet, M., Keller, M. A., Oemer, G., Seppi, D., Haslbeck, M., Pansi, K., Dieplinger, H. & Rupp, B. (2017). Structure 25, 1907–1915.

Naschberger A., Bowler M. W., Rupp B. (2018a). Structural Evidence for a Role of the Multi-functional Human Glycoprotein Afamin in Wnt Transport. European Synchrotron Radiation Facility (ESRF). doi:10.15151/ESRF-DC-142893590

Naschberger A., Bowler M. W., Rupp B. (2018b). Controlled dehydration, structural flexibility, and Gadolinium MRI contrast compound binding in human plasma glycoprotein afamin. European Synchrotron Radiation Facility (ESRF). doi:10.15151/ESRF-DC-142915526

Naschberger A., Bowler M. W., Rupp B. (2019). Gadolinium MRI contrast compound binding in human plasma glycoprotein afamin. European Synchrotron Radiation Facility (ESRF). doi:10.15151/ESRF-DC-186857652

Newman, J. (2006). Acta Cryst. D62, 27–31.

Nile, A. H. & Hannoush, R. N. (2019). J Biol Chem 294, 726–736.

Nurizzo, D., Bowler, M. W., Caserotto, H., Dobias, F., Giraud, T., Surr, J., Guichard, N., Papp, G., Guijarro, M., Mueller-Dieckmann, C., Flot, D., McSweeney, S., Cipriani, F., Theveneau, P. & Leonard, G. A. (2016). Acta Cryst. D72, 966–975.

Nusse, R. & Varmus, H. (2012). EMBO J 31, 2670–2684.

Pellegrini, E., Piano, D. & Bowler, M. W. (2011). Acta Cryst. D67, 902–906.

Raj, I., Sadat Al Hosseini, H., Dioguardi, E., Nishimura, K., Han, L., Villa, A., de Sanctis, D. & Jovine, L. (2017). Cell 169, 1315–1326.e1317.

Rupp, B. (2018). Structure 26, 919–923.

Russo Krauss, I., Sica, F., Mattia, C. A. & Merlino, A. (2012). Int. J. Mol. Sci. 13, 3782.

Sanchez-Weatherby, J., Bowler, M. W., Huet, J., Gobbo, A., Felisaz, F., Lavault, B., Moya, R., Kadlec, J., Ravelli, R. B. G. & Cipriani, F. (2009). Acta Cryst. D65, 1237–1246.

Scherer, M., Klingl, S., Sevvana, M., Otto, V., Schilling, E.-M., Stump, J. D., Müller, R., Reuter, N., Sticht, H., Muller, Y. A. & Stamminger, T. (2014). PLOS Pathogens 10, e1004512.

Schultz, W. W., van Andel, P., Sabelis, I. & Mooyaart, E. (1999). Magnetic resonance imaging of male and female genitals during coitus and female sexual arousal, BMJ 319, 1596–1600.

Seeber, B., Morandell, E., Lunger, F., Wildt, L. & Dieplinger, H. (2014). Reprod. Biol. Endocrin. 12, 88.

Shen, C.-T., Wei, W.-J., Qiu, Z.-L., Song, H.-J. & Luo, Q.-Y. (2016). Mol. Cell. Endocrin. 434, 108–115.

Strong, M., Sawaya, M. R., Wang, S., Phillips, M., Cascio, D. & Eisenberg, D. (2006). Proc. Nat. Acad. Sci. U.S.A. 103, 8060–8065.

Svensson, O., Gilski, M., Nurizzo, D. & Bowler, M. W. (2018). Acta Cryst. D74. 433–440

Svensson, O., Malbet-Monaco, S., Popov, A., Nurizzo, D. & Bowler, M. W. (2015). Acta Cryst. D71, 1757–1767.

Tickle, I. J., Flensburg, C., Keller, P., Paciorek, W., Sharff, A., Vonrhein, C., Bricogne, G. & (http://staraniso.globalphasing.org/cgi-bin/staraniso.cgi). Cambridge, U. K. G. P. L. (2018). Cambridge, United Kingdom: Global Phasing Ltd., (http://staraniso.globalphasing.org/cgibin/staraniso.cgi).

Tickle, I. J., Sharff, A., Flensburg, C., Smart, O. S., Keller, P., Vonrhein, C., Paciorek, W. & Bricogne, G. (2016). http://staraniso.globalphasing.org.

Voegele, A. F., Jerković, L., Wellenzohn, B., Eller, P., Kronenberg, F., Liedl, K. R. & Dieplinger, H. (2002). Biochem. 41, 14532–14538.

Weichenberger, C., Pozharski, E. & Rupp, B. (2017). Acta Cryst. D73, 211–222.

Wheeler, M. J., Russi, S., Bowler, M. G. & Bowler, M. W. (2012). Acta Cryst. F68, 111–114.

Winn, M. D., Ballard, C. C., Cowtan, K. D., Dodson, E. J., Emsley, P., Evans, P. R., Keegan, R. M., Krissinel, E. B., Leslie, A. G. W., McCoy, A., McNicholas, S. J., Murshudov, G. N., Pannu, N. S., Potterton, E. A., Powell, H. R., Read, R. J., Vagin, A. & Wilson, K. S. (2011). Acta Cryst. D67, 235–242.

Wlodawer, A., Dauter, Z., Porebski, P. J., Minor, W., Stanfield, R., Jaskolski, M., Pozharski, E., Weichenberger, C. X. & Rupp, B. (2018). FEBS J 285, 444–466.

Zander, U., Hoffmann, G., Cornaciu, I., Marquette, J. P., Papp, G., Landret, C., Seroul, G., Sinoir, J., Rower, M., Felisaz, F., Rodriguez-Puente, S., Mariaule, V., Murphy, P., Mathieu, M., Cipriani, F. & Marquez, J. A. (2016).). Acta Cryst. D72, 454–466.

Zerrad, L., Merli, A., Schroeder, G. F., Varga, A., Graczer, E., Pernot, P., Round, A., Vas, M. & Bowler, M. W. (2010). J. Biol. Chem. 286, 14040–14048

